# The cellular response towards lanthanum is substrate specific and reveals a novel route for glycerol metabolism in *Pseudomonas putida* KT2440

**DOI:** 10.1101/567529

**Authors:** Matthias Wehrmann, Maxime Toussaint, Jens Pfannstiel, Patrick Billard, Janosch Klebensberger

## Abstract

Ever since the discovery of the first rare earth element (REE)-dependent enzyme, the physiological role of lanthanides has become an emerging field of research due to the potential environmental implications and biotechnological opportunities. In *Pseudomonas putida* KT2440, the two pyrroloquinoline quinone-dependent alcohol dehydrogenases (PQQ-ADHs) PedE and PedH are inversely produced in response to La^3+^-availability. This REE-switch is orchestrated by a complex regulatory network including the PedR2/PedS2 two-component system and is important for efficient growth on several alcoholic volatiles. As *P. putida* is exposed to a broad variety of organic compounds in its natural soil habitat, the cellular responses towards La^3+^ during growth on various carbon and energy sources were investigated with a differential proteomic approach. Apart from the Ca^2+^-dependent enzyme PedE, the differential abundance of most other identified proteins was conditional and revealed a substrate specificity. Concomitant with the proteomic changes, La^3+^ had a beneficial effect on lag-phases while causing reduced growth rates and lower optical densities in stationary phase during growth on glycerol. When these growth phenotypes were evaluated with mutant strains, a novel metabolic route for glycerol utilization was identified that seems to be functional in parallel with the main degradation pathway encoded by the *glpFKRD* operon. The newly discovered route is initiated by PedE and/or PedH, which most likely convert glycerol to glyceraldehyde. In the presence of lanthanum, glyceraldehyde seems to be further oxidized to glycerate, which, upon phosphorylation to glycerate-2-phosphate by the glycerate kinase GarK, is finally channelled into the central metabolism.

**Importance:** The biological role of rare earth elements has long been underestimated and research has mainly focused on methanotrophic bacteria. We have recently demonstrated that *P. putida*, a plant growth promoting bacterium that thrives in the rhizosphere of various feed crops, possesses a REE-dependent alcohol dehydrogenase (PedH), but knowledge about lanthanide-dependent effects on physiological traits in non-methylotrophic bacteria is still scarce. This study demonstrates that the cellular response of *P. putida* KT2440 towards La^3+^ is mostly substrate specific and that during growth on glycerol, La^3+^ has a severe effect on growth parameters. We provide compelling evidence that the observed physiological changes are linked to the catalytic activity of PedH and thereby identify a novel route for glycerol metabolism in this biotechnological relevant organism. Overall, these findings demonstrate that lanthanides can alter important physiological traits of non-methylotrophic bacteria, which might consequently influence their competitiveness during colonization of various environmental niches.

## Introduction

The rhizosphere, defined as the narrow region of soil surrounding plant roots, is one of the most complex ecosystems on earth containing a multitude of organisms from different taxa including fungi, oomycetes, nematodes, protozoa, algae, viruses, archaea and arthropods as well as up to 10^8^ soil dwelling bacteria per gram of fresh root (1–3). Its diversity is mainly shaped by root exudates, a complex mixture of organic compounds including carbohydrates, amino acids, or carbon acids (4, 5) and plant-, fungal-, and bacteria-derived volatiles (VOCs) such as alkenes, alcohols, terpenes or benzenoids (6, 7). As such, it is not surprising that the soil-dwelling organism *P. putida* KT2440 is equipped with a broad diversity of metabolic pathways in order to maximize its cellular fitness in different environmental niches including the rhizosphere (8–10). For efficient growth on various alcoholic VOC substrates, it uses a periplasmic oxidation system consisting of the pyrroloquinoline quinone-dependent alcohol dehydrogenases (PQQ-ADHs) PedE and PedH (11, 12). The enzymes appear to be functionally redundant but differ in their metal cofactor dependency. PedE is a Ca^2+^-dependent enzyme, whereas PedH requires the presence of rare earth elements (REEs) of the lanthanide series (Ln^3+^) for catalytic activity (12, 13).

Although being among the most ubiquitous metals in the earth’s crust, REEs were long considered to be of no biological relevance due to their low solubility under environmental conditions (14). Indeed, the only known and characterized REE-dependent enzymes thus far belong to the family of PQQ-ADHs of methano- and methylotrophic bacteria as well as the non-methylotrophic organism *P. putida* KT2440 (12, 15–20). A characteristic aspartic acid that is additionally present in the metal coordination sphere of these enzymes is associated with Ln^3+^-binding. Notably, this specific amino acid residue has been found in the genome of many bacteria from various origins indicating a broad distribution of Ln^3+^-dependent enzymes (21–25). Very recently, another Ln^3+^-binding protein, called lanmodulin, was identified in *M. extorquens* AM1 (26). This periplasmic protein, which shows structural similarities with the Ca^2+^-binding protein calmodulin, is able to bind up to four Ln^3+^ ions per protein with picomolar affinity and changes its conformation from a largely disordered to a compact, ordered state upon REE binding. Although its exact cellular role has yet to be established, it has been speculated to play a role in Ln^3+^-uptake. Further, homologous genes have only been identified in the genome of some other species of *Methylobacteria* and *Bradyrhizobia*.

In addition to their functional role as metal cofactor, several studies have recently investigated the effect of REEs on cellular physiology in different methano- and methylotrophic organisms (20, 27–32). Some of these studies found different physiological traits to be influenced in response to Ln^3+^-availability including changes in metabolite cross-feeding, growth rates and-yields, or biofilm formation. It is further interesting to note that REEs have been used as micro-fertilizers, especially in China, for over 30 years, as Ln^3+^-supplementation can be associated with increased growth of different food crops including rice, mungbean, maize, and coconut plants (33–37).

The aforementioned results suggest that apart from the inverse transcriptional regulation of PQQ-ADHs, which has been described in detail for different organisms including *P. putida* (12, 19, 27, 38–42), additional responses towards REEs exist and could depend on the specific organism and/or environmental context. To investigate the existence of such conditional cellular responses in the non-methylotrophic organism *P. putida* KT2440, we used a differential proteomic approach during growth on various carbon and energy sources that reflect the metabolic diversity of the rhizosphere. From these experiments, we found that the Ca^2+^-dependent PQQ-ADH PedE was the only protein showing a differential abundance during growth on all carbon and energy sources tested. The vast majority of identified proteins were differentially abundant only under one specific growth condition. During growth on glycerol and 2-phenylethanol, which both represent substrates for PedE and PedH, a disproportionally high number of metabolism related proteins were more abundant in the presence of La^3+^, while this was not the case during growth on citrate and glucose, carbon and energy sources that do not represent substrates for the two PQQ-ADHs. In addition, physiological characteristics, such as growth rates and the lag-phase, of cultures could be linked to the differential activity pattern of PedE and PedH during growth on glycerol. Based on these results, we were able to identify and reconstruct a novel metabolic route for glycerol utilisation, which depends on PedE and/or PedH activity. This route seems to operate in conjunction with the previously described major degradation pathway initiated by the glycerol kinase GlpK and most likely ensures efficient growth of *P. putida* on this polyol substrate.

## Materials and Methods

### Bacterial strains, plasmids and culture conditions

A complete list of all strains, plasmids, and primers used in this study can be found in **Table S1** and **Table S2**. All *Pseudomonas putida* and *Escherichia coli* strains were maintained on solidified LB medium (43). If necessary, 40 µg/mL kanamycin or 20 µg/mL 5-fluorouracil were added for maintenance and/or selection. For growth, liquid LB medium or a modified M9 salt medium (12) supplemented with 5 mM 2-phenylethanol, 25 mM succinate, 10 mM glucose, 10 mM citrate, 20 mM DL-glycerate, or 20 mM glycerol as sole source of carbon and energy was used. If not stated otherwise, precultures were grown overnight in test tubes with 3 ml M9 medium supplemented with succinate at 30°C and 180 rpm. The next day, cells were washed twice with M9 medium without supplemented C-source, and used to inoculate 200 µL of M9 medium supplemented with the desired C-source in a 96-well microtiter plate (Falcon, product no. 353047 or Sarstedt, product no. 83.3924) and incubated at 30°C and 250 rpm in a microplate reader (Xenius, Safas Monaco) or 28°C and 220 rpm in a rotary shaker (Forma, Thermo Scientific). Maximum growth rates (*µ*_max_) and lag-times (*λ*) were estimated based on fitting the natural logarithm of the relative OD_600_ values (ln(N/N_0_), with N being OD_600_ at time t) with the Richards growth model using the “grofit” package in R (44). As OD_600_ decreased directly upon begin of the experiment, ln(N/N_t = 3 h_) was used instead of ln(N/N_0_) for better fit. Differences in lag-times, growth rates, and maximal OD_600_ during stationary phase (OD_600_^max^) were evaluated by statistical analysis in GraphPad PRISM using a two-tailed *t*-test (*α* = 0.05, *n* = 3).

### Construction of plasmids

Deletion plasmids pJOE-calA, pJOE-garK, pJOE-glp, and pMW08 were constructed as follows: the 650-bp to 1000-bp regions upstream and downstream of the *calA* (PP_2426), *garK* (PP_3178), *glpFKRD* (PP_1076 to PP_1973), or *glcDEF* (PP_3745 to PP_3747) genes were amplified from genomic DNA of *P. putida* KT2440 using primers PcalA1 to PcalA4, PgarK1 to PgarK4, Pglp1 to Pglp4, or MWH03 to MWH06 (**Table S2**). The two up- and downstream fragments and BamHI-digested pJOE6261.2 were then joined together using one-step isothermal assembly (45).

### Strain constructions

Deletion mutant strains were constructed as previously described (46). Briefly, the integration vector pJOE6261.2 harbouring the up- and downstream regions of the target gene(s) was transformed into *P. putida* KT2440 *Δupp* (KT2440*). Kanamycin (Kan) resistant and 5-fluorouracil (5-FU) sensitive clones were selected and one of these was incubated in LB medium at 30°C for 24 h. The cell suspension was then plated on M9 minimal agar plates containing 25 mM succinate and 20 µg ml^-1^ 5-FU. Clones that carried the desired gene deletion were identified by colony PCR of the 5-FU^r^ Kan^s^ clones using primer pair PcalA1/PcalA4, PgarK1/PgarK4, Pglp1/Pglp4, or MWH03/MWH06.

### Protein extraction for comparative proteome analysis

For comparative proteome analysis experiments, 50 ml M9 medium supplemented with citrate, glucose, glycerol or 2-phenylethanol and 0 or 10 µM LaCl_3_ were inoculated with an OD_600_ of 0.05 from succinate precultures of strain *P. putida* KT2440* in 250 ml polycarbonate Erlenmeyer flasks and incubated at 30°C and 180 rpm. When cell cultures reached an OD_600_ of > 0.4, cells were harvested by centrifugation for 15 min at 6000 x g and 4°C. Cell pellets were resuspended in 1 ml sample buffer (150 mM Tris-HCl pH 6.8; 2 % SDS; 20 mM dithiothreitol) and heated for 5 min at 95°C with gentle shaking. Subsequently, samples were centrifuged for 15 min at 21000 x g and 4°C, and the supernatants were stored in new reaction tubes at −20 °C. In a next step, proteins were precipitated using chloroform-methanol (47) and pellets were resuspended in Tris-buffered (50 mM, pH 8.5) urea (6 M). Protein concentrations were determined by the Bradford assay (48).

### In-solution digest of proteins and peptide purification with C18 Stage Tips

To 25 µg protein in 60 µl Tris-buffered (50 mM, pH 8.5) urea (6 M), DTT was added to a final concentration of 10 mM to guarantee reduction of cysteines. Samples were incubated for 30 min at 56 °C under shaking at 1000 rpm. Alkylation of cysteines was performed by adding 30 mM iodoacetamide and incubation for 45 min at room temperature in the dark. Alkylation was stopped by adding 50 mM DTT and samples were incubated for another 10 min at RT. 500 ng LysC protease (Roche) in 50 mM Tris buffer (pH 8.5) was added and samples were digested overnight at 30 °C. Next, the urea in the reaction mixture was diluted to 2 M by adding the appropriate amount of Tris buffer (50 mM, pH 8.5). 1 µg trypsin (Roche) in Tris buffer (50 mM, pH 8.5) was added and digestion was continued for 4 hours at 37 °C. The digest was stopped by addition of 3 µl 10% (v/v) trifluoroacetic acid (TFA). Next, peptide mixtures were concentrated and desalted on C18 stage tips (49) and dried under vacuum. Samples were dissolved in 20 µl 0.1% (v/v) TFA. Aliquots of 1 µl were subjected to nanoLC-MS/MS analysis.

### Mass spectrometry analysis

NanoLC-ESI-MS/MS experiments were performed on an EASY-nLC 1200 system (Thermo Fisher Scientific) coupled to a Q-Exactive Plus mass spectrometer (Thermo Fisher Scientific) using an EASY-Spray nanoelectrospray ion source (Thermo Fisher Scientific). Tryptic peptides were directly injected to an EASY-Spray analytical column (2 μm, 100 Å PepMapRSLC C18, 25 cm × 75 μm, Thermo Fisher Scientific) operated at constant temperature of 35 °C. Peptides were separated at a flow rate of 250 nL/min using a 240 min gradient with the following profile: 2% - 10% solvent B in 100 min, 10% - 22% solvent B in 80 min, 22% - 45% solvent B in 55 min, 45% - 95% solvent B in 5 min and isocratic at 90% solvent B for 15 min. Solvents used were 0.5 % acetic acid (solvent A) and 0.5% acetic acid in acetonitrile/H2O (80/20, v/v, solvent B). The Q Exactive Plus was operated under the control of XCalibur 3.0.63 software. MS spectra (m/z = 300-1600) were detected in the Orbitrap at a resolution of 70000 (m/z = 200) using a maximum injection time (MIT) of 100 ms and an automatic gain control (AGC) value of 1 x 10^6^. Internal calibration of the Orbitrap analyzer was performed using lock-mass ions from ambient air as described elsewhere (50). Data dependent MS/MS spectra were generated for the 10 most abundant peptide precursors in the Orbitrap using high energy collision dissociation (HCD) fragmentation at a resolution of 17500, a normalized collision energy of 27 and an intensity threshold of 1.3 x 10^5^. Only ions with charge states from +2 to +5 were selected for fragmentation using an isolation width of 1.6 Da. For each MS/MS scan, the AGC was set at 5 x 10^5^ and the MIT was 100 ms. Fragmented precursor ions were dynamically excluded for 30 s within a 5 ppm mass window to avoid repeated fragmentation.

### Protein quantification and data analysis

Raw files were imported into MaxQuant (51) version 1.6.0.1 for protein identification and label-free quantification (LFQ) of proteins. Protein identification in MaxQuant was performed using the database search engine Andromeda (52). MS spectra and MS/MS spectra were searched against *P. putida* KT2440 protein sequence database downloaded from UniProt (53). Reversed sequences as decoy database and common contaminant sequences were added automatically by MaxQuant. Mass tolerances of 4.5 ppm (parts per million) for MS spectra and 20 ppm for MS/MS spectra were used. Trypsin was specified as enzyme and two missed cleavages were allowed. Carbamidomethylation of cysteines was set as a fixed modification and protein N-terminal acetylation and oxidation were allowed as variable modifications. The ‘match between runs’ feature of MaxQuant was enabled with a match time window of one minute and an alignment time window of 20 minutes. Peptide false discovery rate (FDR) and protein FDR thresholds were set to 0.01.

Statistical analysis including *t*-tests and principal component analysis (PCA) were performed using Perseus software version 1.6.0.2 (54). Matches to contaminant (e.g., keratins, trypsin) and reverse databases identified by MaxQuant were excluded from further analysis. Proteins were considered for LFQ (label free quantification) if they were identified by at least two peptides. First, normalized LFQ values from MaxQuant were log2 transformed. Missing values were replaced from normal distribution using a width of 0.2 and a downshift of 2.0. Statistical differences between two sample groups were determined using an unpaired *t*-test and a *p*-value < 0.01 and a regulation factor > 2 (log2 fold-change > 1) were considered as significant change in protein abundance. The mass spectrometry proteomics data will be deposited to the ProteomeXchange Consortium via the PRIDE (55) partner repository (submitted).

### Purification and activity measurement of PQQ-ADHs PedE and PedH

To measure the activity of the two PQQ-ADHs PedE and PedH, the enzymes were expressed in *E. coli* BL21(DE3) cells using plasmids pMW09 and pMW10, and purified by affinity chromatography as described elsewhere (12). The activities with the four substrates 2-phenylethanol, citrate, glucose and glycerol were determined at a concentration of 10 mM using a previously described colorimetric assay (12) with one minor modification. To represent the growth conditions, 1 µM La^3+^ instead of 1 µM Pr^3+^ was used as metal cofactor for PedH.

## Acknowledgements

The authors would like to thank Prof. Bernhard Hauer for his continuous support. The authors further declare that the research was conducted in the absence of any commercial or financial relationships that could be construed as a potential conflict of interest.

## Results

We have recently demonstrated that the two PQQ-ADHs PedE and PedH are inversely regulated dependent on the presence of rare earth elements (REEs) and that a complex signalling network, which includes the activity of the PedR2/PedS2 two-component system, orchestrates this regulation (12, 38). To identify whether a global cellular response of *P. putida* KT2440 towards REEs beyond the regulation of the PQQ-ADHs exists, we used a comparative proteomic analysis during growth on four different carbon and energy sources, namely 2-phenylethanol, glycerol, glucose, and citrate.

### Evaluation of proteomics data

Proteins were extracted from cells of *P. putida* by SDS to enable extraction of cytoplasmic as well as transmembrane proteins followed by label free nano-LC-MS/MS quantification. In total, 2771 proteins with at least two unique peptides and an FDR ≤ 1% were identified and quantified by our proteomics approach, corresponding to approximately 50% of the *P. putida* KT2440 proteome. Principal component analysis revealed high reproducibility for sample replicates and distinct patterns for the different carbon sources (**Fig. S1**). The majority of proteins was increased or decreased in abundance in response to the different carbon and energy sources. In contrast, minor differences were observed in the presence or absence of La^3+^ during growth on the same carbon and energy source. Proteins that exhibited a 2-fold or higher change in abundance between different growth conditions and a *p*-value ≤ 0.01 were considered as differentially abundant.

### Effect of lanthanum on protein abundance during growth with different substrates

According to the aforementioned criteria, 56 proteins were identified as differentially abundant comparing growing cells of *P. putida* in the presence and absence of La^3+^ with different carbon sources (**Fig. 1, Table 3, Table S3-S5**). In these studies, only the Ca^2+^-dependent PQQ-ADH PedE (PP_2674) showed a decreased abundance in response to La^3+^ during growth on all four different carbon and energy sources. The Ln^3+^-dependent PQQ-ADH PedH (PP_2679) showed an increased abundance in response to La^3+^ during growth on glucose, glycerol, and 2-phenylethanol, whereas an uncharacterized pentapeptide repeat containing protein (PP_2673) that is directly upstream of *pedE* showed a decreased abundance during growth on glycerol and 2-phenylethanol (**Fig. 1**). The remaining 53 proteins were only identified under one specific growth condition (**Table 3, Table S3-S5**).

**Figure 1:**
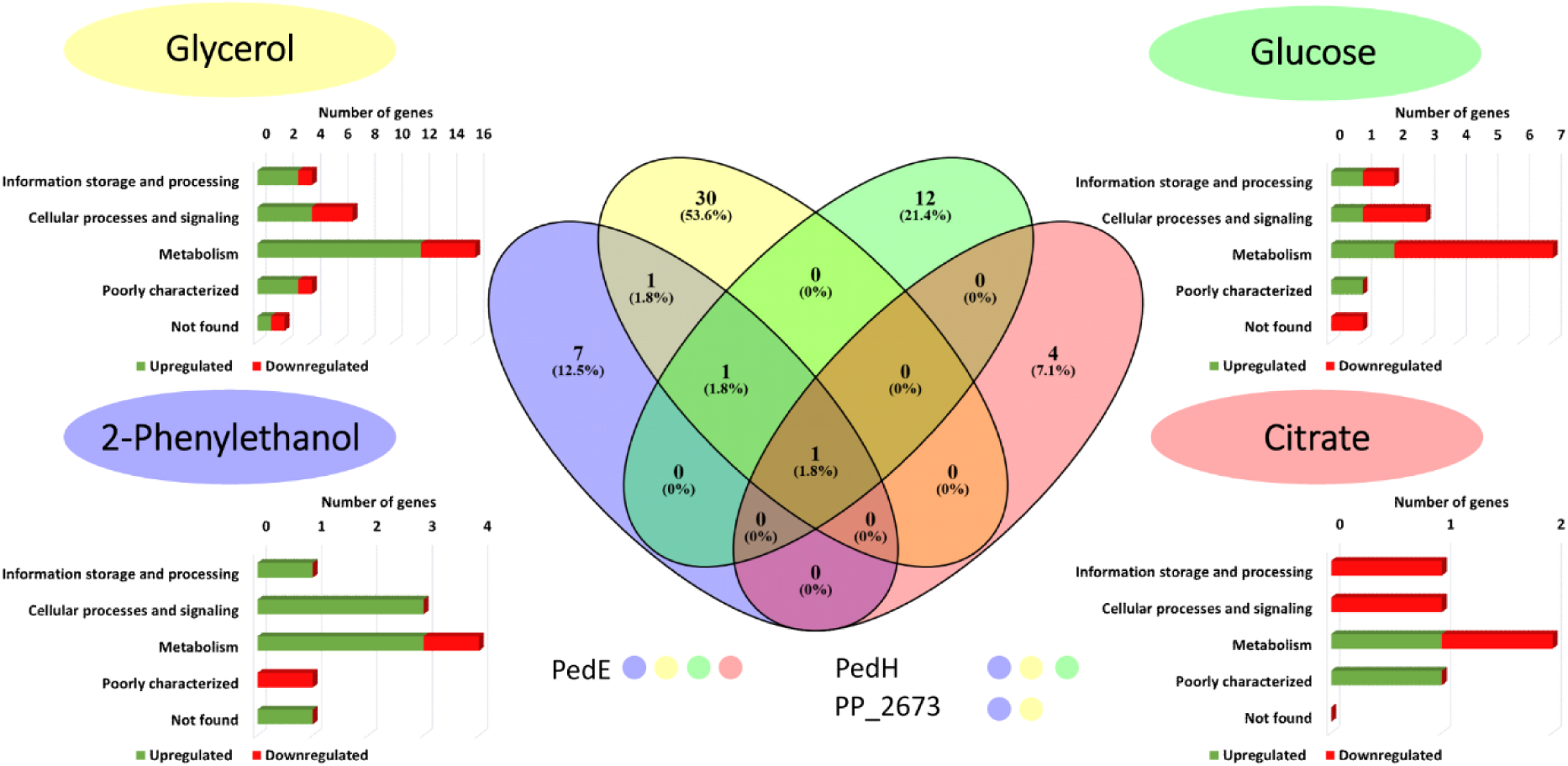
Venn-diagram (middle) of differentially produced proteins in response to 10 µM La^3+^ during growth with glycerol, glucose, 2-phenylethanol and citrate. Proteins that showed up under several growth conditions are stated under the diagram with colour code for classification (yellow dot = glycerol; green dot = glucose; blue dot = 2-phenylethanol; red dot = citrate). Classifications of differentially expressed proteins according to the Cluster of Orthologous Groups database are depicted for each substrate.

During growth on 2-phenylethanol and glycerol, a majority of the identified proteins was increased in abundance (80% and 70%) in response to La^3+^ (**Table 3, Table S3**). For glucose and citrate this was different, as most of the identified proteins were found to be less abundant in response to La^3+^ (36% and 40% during growth on glucose and citrate; **Table S4 and Table S5**). Notably, the majority of the identified proteins were related to metabolism according to the cluster of orthologous protein groups (COG) database (56). To test whether the observed conditional proteomic response is linked to PedE and/or PedH activity, we determined the corresponding enzyme activities with all four carbon and energy sources (**Table 1**). Apart from the already known substrate 2-phenylethanol, PedE and PedH also showed activity with glycerol, whereas no activity could be detected with citrate or glucose.

**Table 1:**
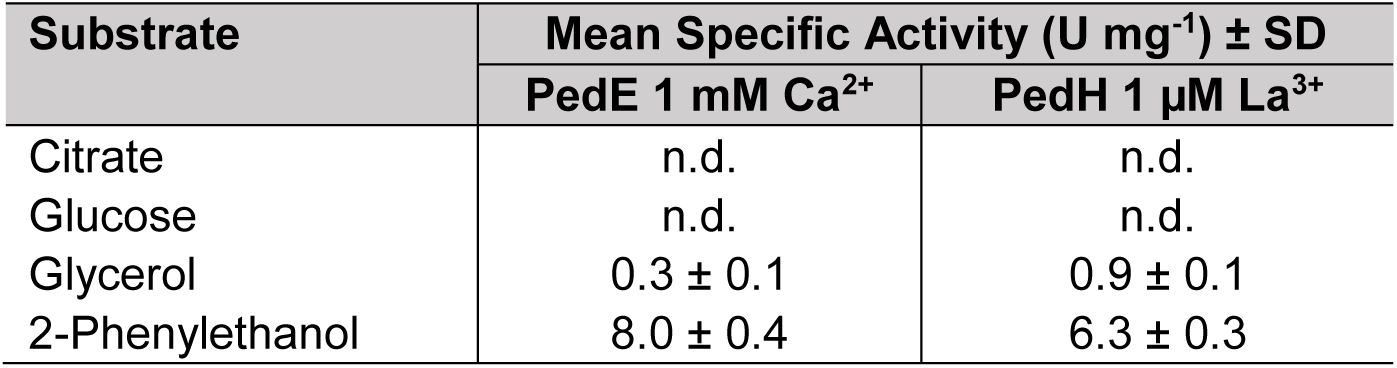
Specific enzyme activities of purified PedE and PedH with the four tested growth substrates at 10 mM measured with 2,6-dichlorophenolindophenol (DCPIP) dependent colorimetric assay. Data represent the average of biological triplicates with according standard deviation. Activities below detection limit are indicated (n. d.).

### Effect of lanthanum during growth on glycerol

From our proteomic- and biochemical data, we speculated that PedE and PedH activity could play a beneficial role during glycerol metabolism of *P. putida* KT2440. As the degradation pathway and growth characteristics of this organism have been recently characterized in great detail (57, 58), we wanted to have a closer look on the effect of La^3+^ during growth on this specific carbon and energy source. In these experiments (**Fig. 2A, Table 2**), we consistently observed a shorter lag-phase (*λ*) of the cultures in response to La^3+^-availability (10.2 ± 0.2 h vs. 17.3 ± 0.2). Additionally, the corresponding values of the specific growth rates (*μ*_max_, 0.201 ± 0.004 vs. 0.341 ± 0.010 h^-1^) and the maximal OD_600_ in stationary phase (OD_600_^max^; 0.680 ± 0.014 vs. 0.884 ± 0.004) of the cultures differed in the presence or absence of La^3+^, respectively. As the purified PedH enzyme showed a 3-fold higher specific activity towards glycerol compared to PedE *in vitro* (0.9 ± 0.1 U mg^-1^ vs. 0.3 ± 0.1 U mg^-1^; **Table 1**), we speculated that this increased glycerol conversion by PedH could be the underlying cause for the observed differences in growth parameters. When subsequently a Δ*pedE* Δ*pedH* strain was analysed for growth in the presence and absence of La^3+^ (**Fig. 2B, Table 2**), no significant differences in *λ* and μ_max_ were observed for the Δ*pedE* Δ*pedH* strain in response to La^3+^ while, although less profound, small differences in OD_600_^max^ were still detected. Further, under both conditions the double deletion strain showed a lag-phase that was undistinguishable from that of the parental strain in the absence of La^3+^ but dramatically longer than that of the parental strain in the presence of La^3+^ (17.5 ± 0.3 h in the absence of La^3+^ and 17.2 ± 0.4 h in the presence of 10 μM La^3+^). Interestingly, the growth rates under both conditions (0.276 ± 0.008 h^-1^ and 0.291 ± 0.012 h^-1^) were significantly higher (*p* < 0.01) than those of the parental strain in presence of La^3+^ while still being significantly below (*p* < 0.05) those of the parental strain in the absence of La^3+^.

**Figure 2:**
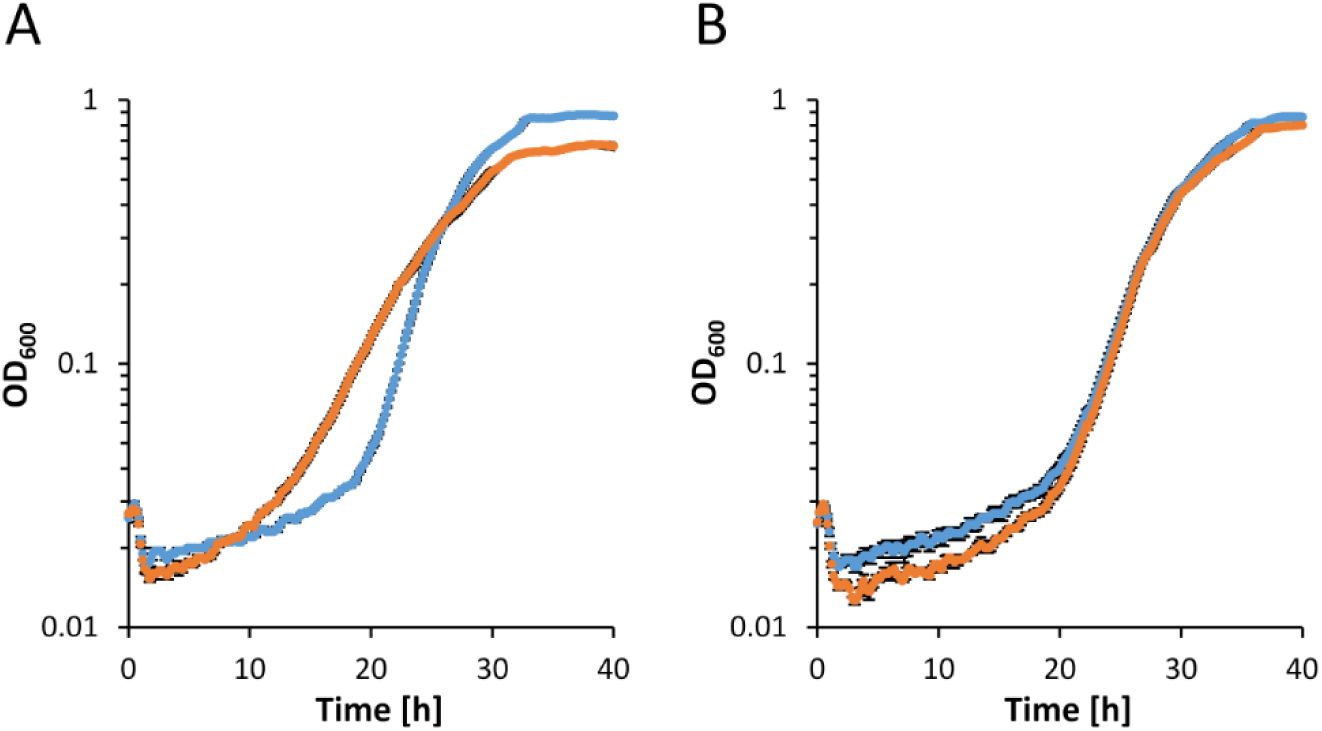
Growth of strains (**A**) KT2440* and (**B**) Δ*pedE* Δ*pedH* in M9 minimal medium supplemented with 20 mM glycerol in the absence (blue dots) or presence of 10 μM La^3+^ (orange dots) in 96-well microtiter plates at 30°C and 250 rpm. Data represent average of biological triplicates with corresponding standard deviation.

**Table 2:** Lag-times (*λ*), maximal OD_600_ during stationary phase (OD_600_^max^), and maximal growth rates (*μ*_max_) of different *P. putida* strains during growth with M9 medium supplemented with 20 mM glycerol and 0 μM or 10 μM La^3+^ incubated in microtiter plates at 30°C and 250 rpm (see also Fig. 2 and Fig.5). Maximal growth rates and lag-times were determined by fitting growth curves to the Richards model using “grofit” package in R (44). Cultures were incubated at 250 rpm and 30°C in microplate reader with constant OD_600_ measurement. Data points represent average of biological triplicates with corresponding error (*λ, μ*_max_) or standard deviation (OD_600_^max^). No growth parameters could be determined for cultures that did not reach stationary phase during incubation time (n. d.).

| Strain | $\lambda$ [h] | | OD<sub>600</sub><sup>max</sup> | | $\mu_{\max}$ [h<sup>-1</sup>] | |
| --- | --- | --- | --- | --- | --- | --- |
| | 0 μM La<sup>3+</sup> | 10 μM La<sup>3+</sup> | 0 μM La<sup>3+</sup> | 10 μM La<sup>3+</sup> | 0 μM La<sup>3+</sup> | 10 μM La<sup>3+</sup> |
| KT2440* | 17.3 ± 0.2 | 10.2 ± 0.2 | 0.884 ± 0.004 | 0.680 ± 0.014 | 0.341 ± | 0.201 ± 0.004 |
| $\Delta pedE$ | 17.5 ± 0.3 | 17.2 ± 0.4 | 0.869 ± 0.002 | 0.802 ± 0.013 | 0.276 ± | 0.291 ± 0.012 |
| $\Delta garK$ | 17.1 ± 0.3 | n.d. | 0.805 ± 0.005 | n.d. | 0.321 ± | n.d. |
| $\Delta calA$ | 16.0 ± 0.3 | 11.1 ± 0.3 | 0.890 ± 0.004 | 0.753 ± 0.006 | 0.336 ± | 0.271 ± 0.008 |
| $\Delta glcDEF$ | 17.0 ± 0.2 | 10.1 ± 0.2 | 0.862 ± 0.016 | 0.678 ± 0.004 | 0.315 ± | 0.207 ± 0.004 |

**Table 3:**
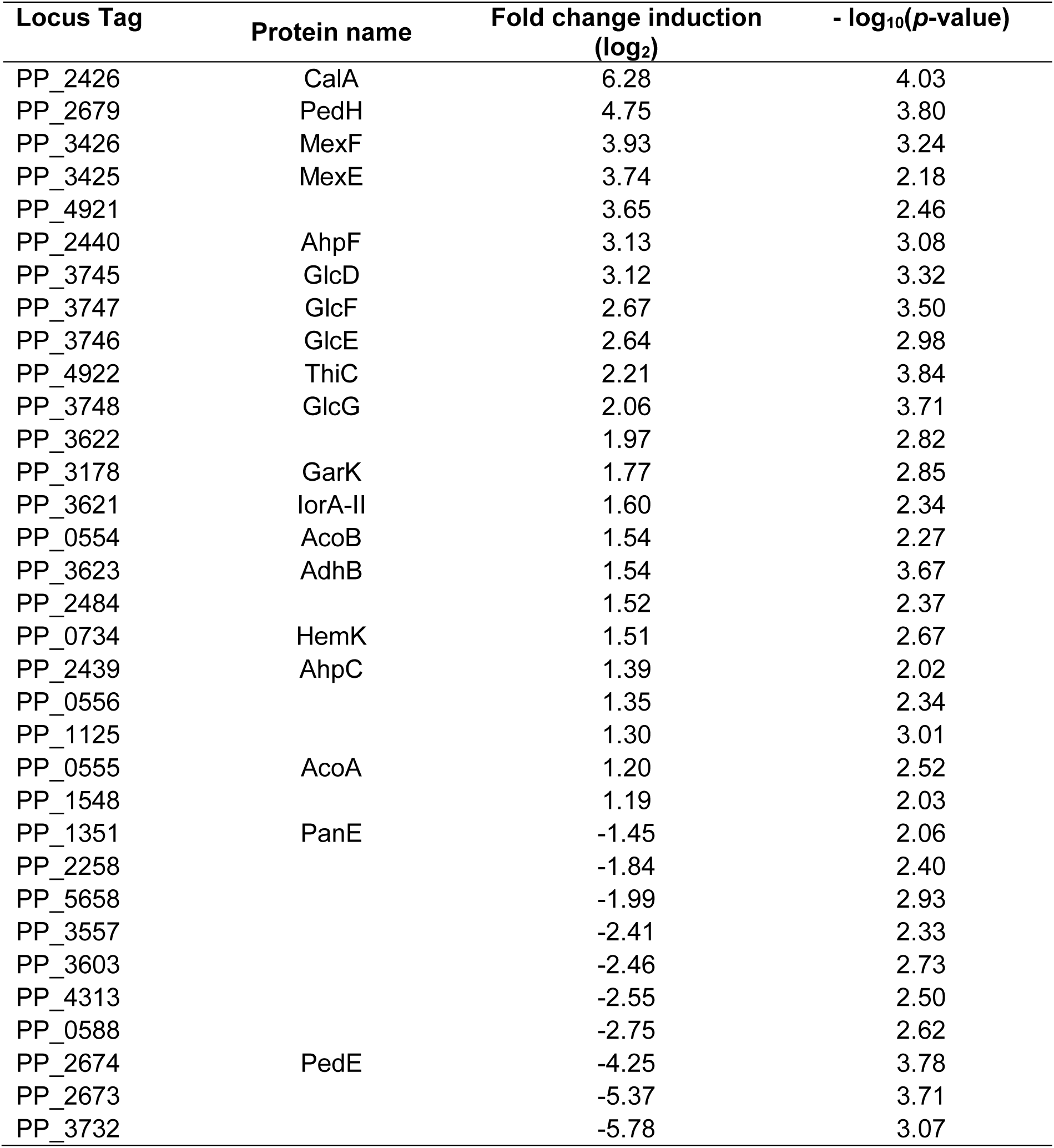
List of regulated proteins in presence of 10 μM La^3+^ compared to the absence of La^3+^ when grown with glycerol as sole C-source.

These results implied that the two PQQ-ADHs can indeed be beneficial for growth on glycerol and that a functionally active PedH enzyme is the underlying cause for the La^3+^-dependent differences in lag-times and growth rates and to some extent also for differences in OD_600_^max^ of KT2440 cultures. As PedE and PedH as well as the remaining proteins that were found to be differentially abundant in response to La^3+^ during growth on glycerol, are not part of the described degradation pathway in *P. putida* KT2440 (57, 58), we hypothesized that an additional metabolic route exists (**Fig. 3**). Based on our proteomic data, this route could be initiated by the activity of PedH and the oxidation of glycerol to glycolaldehyde. In the next steps glycolaldehyde could be oxidized to glycerate by PedH, the aldehyde dehydrogenase AldB-II, or the aldehyde oxidase complex composed of proteins PP_3621 (IorA-II), PP_3622 and PP_3623 (AdhB). After phosphorylation by the glycerate kinase GarK, glycerate-2-phosphate could eventually enter the central metabolism.

**Figure 3:**
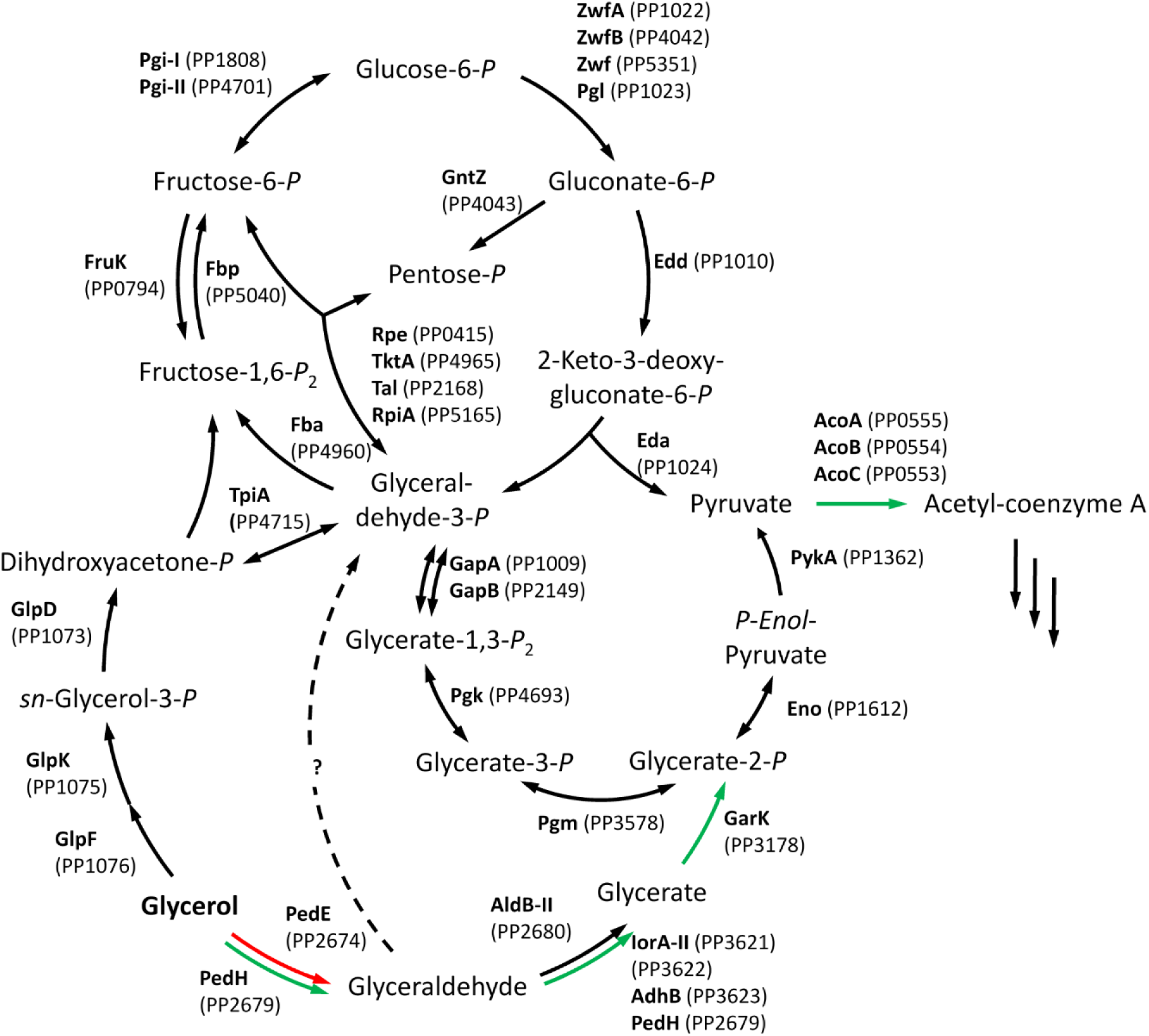
Proteins of the upstream central carbon metabolism of *P. putida* KT2440 including the proposed glycerol degradation pathway via glycerate and the additional hypothetical route through phosphorylation of glyceraldehyde (dotted line). Anticipated metabolic flux by the proteins that were identified as differentially abundant in response to 10 μM La^3+^ during growth on glycerol are colour-coded (green = increased, red = decreased, black = not affected). The figure is inspired by a scheme originally published by Nikel *et al.* (58) and was adapted to include the novel metabolic route(s) identified in this study.

If such a metabolic route exists, a Δ*glpFKRD* deletion strain should still be able to grow with glycerol as sole source of carbon and energy, whereas a Δ*pedE* Δ*pedH* Δ*glpFKRD* deletion mutant should not. To test this scenario, the corresponding strains were constructed and characterized for their growth phenotypes (**Fig. 4A**). We found that *P. putida* KT2440 indeed grew on glycerol independent of the GlpFKRD pathway, although growth was dramatically impaired compared to the parental strain or strain Δ*pedE* Δ*pedH*. When PedE and PedH were additionally deleted, no growth was observed even after a prolonged incubation time of 5 d. This supported our hypothesis that a metabolic route for glycerol next to the GlpFKRD pathway exists and that this route is initiated by PedE and PedH, most likely by the oxidation of glycerol to glyceraldehyde (**Fig. 3**). Given that the route further proceeds *via* glycerate and glycerate-2-phosphate, different cellular concentrations of these metabolites would be expected during growth on glycerol in a mutant that is not able to use the GlpFKRD pathway. Interestingly, a companion study to this work employed a metabolome analysis using glycerol-growing cells of *P. putida* KT2440 and a Δ*glpK* deletion strain, which can only use the proposed novel route *via* PedE and PedH (59). In their experiments, the authors observed that the glycerate concentration measured for the Δ*glpK* strain was dramatically increased compared to the wild type, whereas concentrations of glyceraldehyde and glyceraldehyde-3-phosphate were in the same range for both strains. These data suggested that glycerate is indeed an intermediate during *glpFKRD*-independent growth, and that the activity of downstream proteins represent the bottleneck of the metabolic route leading to the observed accumulation. As our proteomic data indicated the involvement of the predicted glycerate kinase GarK, we constructed a Δ*garK* deletion strain and speculated that this strain should lack the ability to phosphorylate glycerate and would hence be incapable of channelling glycerate-2-phosphate into the central metabolism. We indeed observed no growth of a Δ*garK* mutant on glycerate even after incubation of up to 5 d, while strain Δ*pedE* Δ*pedH* Δ*glpFKRD* could grow and reached OD_600_^max^ within 72 h of incubation under the condition tested (**Fig. 4B**).

**Figure 4:**
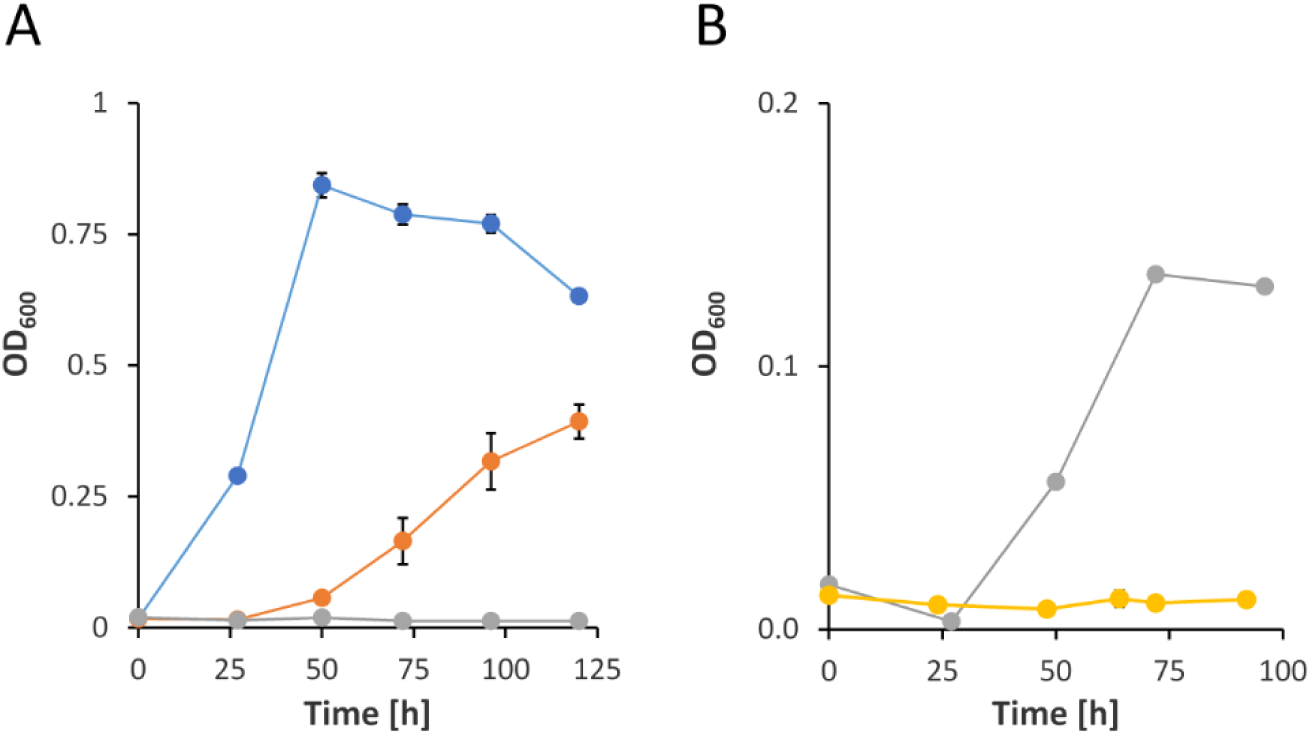
(**A**) Growth of strains Δ*pedE* Δ*pedH* (blue dots), Δ*glpFKRD* (orange dots), and Δ*pedE* Δ*pedH* Δ*glpFKRD* (grey dots) in M9 minimal medium supplemented with 20 mM glycerol in 96-well microtiter plates. (**B**) Growth of strains Δ*pedE* Δ*pedH* Δ*glpFKRD* (grey dots) and Δ*garK* (yellow dots), in M9 minimal medium supplemented with 20 mM DL-glycerate incubated in 96-well microtiter plates at 28°C and 220 rpm. Data represent average of biological triplicates with corresponding standard deviation.

When grown on glycerol, no significant effect on growth rates and lag-times as well as only a minor, but significant, negative effect on the OD_600_^max^ (*p* < 0.01) was observed for the Δ*garK* deletion in the absence of La^3+^. In contrast, the same deletion caused a dramatic growth impairment in the presence of La^3+^ (**Fig. 5A**, **Table 2**) and consequently the stationary phase was not reached during the 40 h of incubation. Hence, no reliable growth parameter could be deduced from these data. It is however obvious that the growth rate was far below the one of the parental strain in the presence of La^3+^.

**Figure 5:**
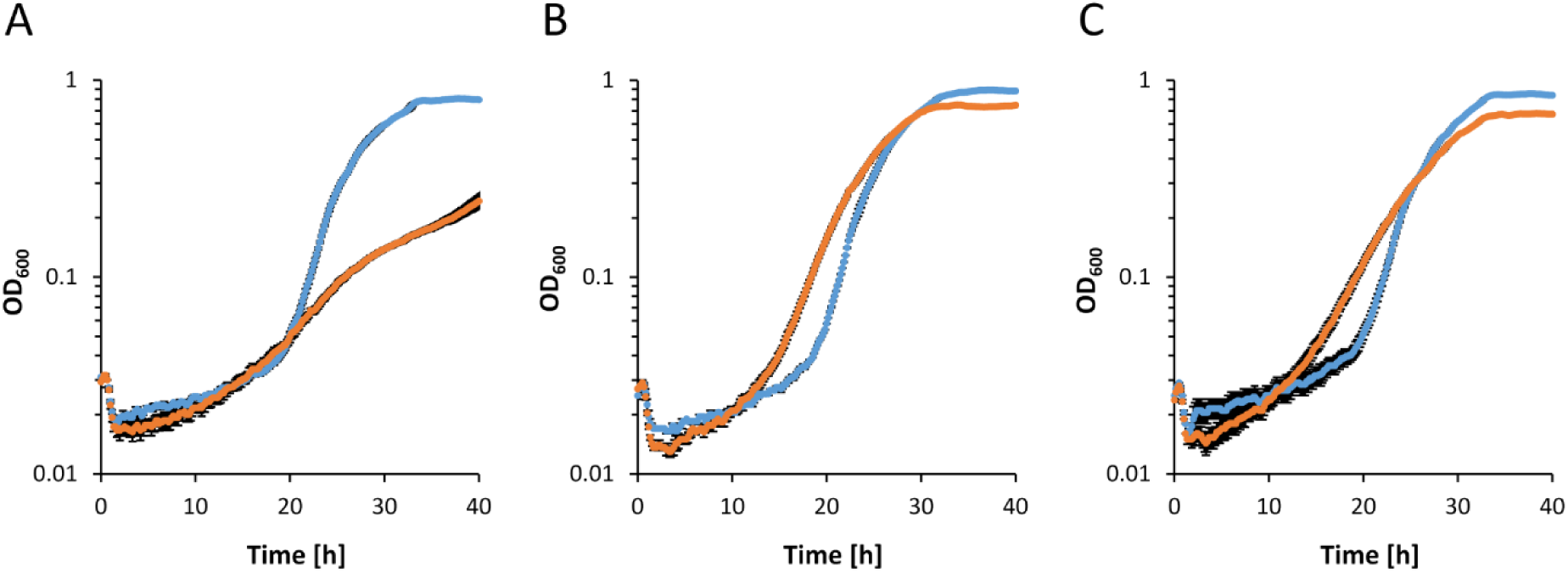
Growth of strains (**A**) Δ*garK*, (**B**) Δ*calA*, and (**C**) Δ*glcDEF* in M9 minimal medium supplemented with 20 mM glycerol in absence (blue dots) or presence 10 μM La^3+^ (orange dots) incubated in 96-well microtiter plates at 30°C and 250 rpm. Data represent average of biological triplicates with corresponding standard deviation.

Notably, some of the most severely upregulated proteins in response to La^3+^ are either related to stress, namely the multidrug efflux pump MexEF and the alkylhydroperoxide reductase subunits AhpC and AhpF, or are enzymes that play no obvious roles within the proposed metabolic route such as CalA, a predicted coniferyl alcohol dehydrogenase, and the glycolate oxidase GlcDEF. To investigate the potential influence of the latter two enzymes, we constructed and analysed the corresponding Δ*calA* and Δ*glcDEF* mutants and tested their growth pattern with glycerol (**Fig. 5B; Table 2**). The Δ*glcDEF* mutant showed a growth behaviour similar to the parental strain. In contrast, the Δ*calA* strain exhibited a significantly increased growth rate (*p* < 0.01) and higher OD_600_^max^ (*p* < 0.01) than the parental strain in the presence of La^3+^ while showing no significant differences in OD_600_^max^ and maximal growth rate and only minor differences (*p* < 0.05) in lag-times in the absence of La^3+^ (**Fig. 5C; Table 2**).

## Discussion

In the present study, the cellular responses of *P. putida* KT2440 towards La^3+^-availability during growth on several carbon and energy sources were investigated. The only protein that showed a differential abundance independent of the substrate used for growth was the Ca^2+^-dependent PQQ-ADH PedE. This result is in line with data from a previous study (38), which demonstrated that the La^3+^-induced downregulation of *pedE* is dependent on the PedS2/PedR2 two-component system that, based on our current observation, seems to be functional under all tested conditions. The other two proteins that showed differential abundance under more than one culture condition (PedH, PP_2673) are both also part of the *ped* gene cluster. Notably, the carbon and energy sources under which these proteins were identified either represent substrates of PedE and PedH, or can be converted by an enzyme that depends on the same PQQ-cofactor, namely the glucose dehydrogenase Gcd. The remaining 53 proteins that showed differential abundance in response to La^3+^ were identified only during growth on one specific carbon and energy source, suggesting a conditional regulation. For glycerol, we provide striking evidence that the increased activity of PedH compared to PedE is the primary cause for the observed proteomic and physiological changes during growth.

Thus far, the degradation of glycerol was described to start by the uptake *via* GlpF, phosphorylation by GlpK, and subsequent GlpD-catalysed oxidation of glycerol-3-phosphate to dihydroxyacetone-3-phosphate (58). In a next step, dihydroxyacetone-3-phosphate is interconverted to glyceraldehyde-3-phosphate and enters the central metabolism. This pathway is negatively regulated by the transcriptional regulator GlpR, and the de-repression of the *glpFKRD* operon is believed to depend on the intracellular concentration of glycerol-3-phosphate, which finally impacts the lag-phase of cultures (57). As such, it was interesting to find that growth on glycerol in the presence of La^3+^ led to a shorter lag phase and lower growth rate of the parent strain, and that a Δ*pedE* Δ*pedH* mutant showed a lag-phase similar to the parent in absence of La^3+^ without any beneficial effect of La^3+^ while still growing with a higher growth rate than the parent strain in presence of La^3+^. Further experiments revealed that a Δ*glpFKRD* deletion strain is still able to grow on glycerol, while a Δ*pedE* Δ*pedH* Δ*glpFKRD* is not. Together with the notion that a Δ*garK* mutant cannot utilize glycerate, this strongly indicates the existence of a novel route for glycerol metabolism, in which PedE and PedH catalyse the initial oxidation of glycerol to glyceraldehyde. In the presence of La^3+^, the route seems to proceed *via* a second oxidation step to glycerate, which is subsequently converted to glycerate-2-phosphate by the activity of GarK (**Fig. 3**). The PedE/PedH-dependent route, despite being important for efficient growth, clearly is not the main route for glycerol metabolism, as the effect of the Δ*pedE* Δ*pedH* deletion on the lag-phase with glycerol is far less severe than deletion of the *glpFKRD* gene cluster. It also appears that the PedE/PedH-dependent route is less efficient than the GlpFKRD pathway, as the overall growth of the Δ*glpFKRD* strain is substantially impaired in comparison to the Δ*pedE* Δ*pedH* strain.

A possible explanation could be the formation of the toxic intermediate glyceraldehyde, which is known for its protein crosslinking properties and the formation of superoxide radicals due to auto-oxidation (60, 61). The observed differences in growth rates and OD_600_^max^ in response to La^3+^ in the parent strain could thus reflect the increased metabolic flux towards glyceraldehyde due to the higher specific activity of PedH compared to PedE. This would also explain the severe La^3+^-dependent growth impairment of the Δ*garK* mutant, as one can assume that even higher concentrations of glyceraldehyde accumulate in a mutant that cannot process glycerate. The notion that the MexEF RND-type transporter proteins, which are involved in efflux of various toxic compounds (62), and the alkylhydroperoxide reductase subunits AhpC and AhpF, which have been linked to ROS detoxification in *P. putida* (63), were also more abundant in presence of La^3+^ during growth on glycerol are supportive of such a hypothesis.

To explain the impact of La^3+^ on the lag-times of cultures, one could speculate that in addition to glycerol-3-phosphate, also other phosphorylated derivatives, such as glycerate-2-phosphate, are able to relieve the repression of *glpFKRD* by GlpR. However, as in the absence of La^3+^ the growth phenotype of the Δ*garK* mutant is indistinguishable from that of the parental strain, and since the growth rate of the parent strain in absence of La^3+^ is still significantly higher than the growth rate of strain Δ*pedE* Δ*pedH*, we postulate that yet another metabolic route is present that contributes to growth without affecting the lag-phase. This second route could proceed *via* the phosphorylation of glyceraldehyde to glyceraldehyde-3-phosphate by the activity of a so-far unknown kinase. Whether both alternative routes to the GlpFKRD pathway are functional in parallel or whether the metabolic flux *via* glycerate is exclusively induced in the presence of La^3+^ is currently unknown and would need to be tested in future studies. Similarly, the question why proteins that cannot be associated to the newly discovered routes for glycerol are among the most differentially abundant proteins in response to La^3+^ remains to be elucidated. It is however worthwhile noting that CalA and GlcDEF are either known (GlcDEF) or predicted (CalA) by the PROSITE software tool (https://prosite.expasy.org/) (64) to be catalytically active on 2-hydroxy acids. As such, potential activities towards pathway intermediates such as glycerate cannot be excluded at the moment.

From our data, the La^3+^-dependent proteomic and physiological changes during growth on glycerol can be explained by a shift in metabolic flux resulting from the differences in specific catalytic activities between PedH and PedE. A similar metabolic-driven interpretation can also be used to explain the proteomic differences during growth on other carbon and energy sources that are known to be substrates for PedE and PedH such as 2-phenylethanol. However, this logic fails to explain the differences observed during growth on glucose and citrate, as they do not represent substrates for PedE and/or PedH. Despite the fact that we currently do not know the underlying cause for the conditional proteomic changes under these conditions, it indicates the presence of additional effects of REEs beside the interaction with PedH and PedS2/PedR2. Such effects could include the inhibition of protein functions by mismetallation (65, 66), changes in the physiology of the outer membrane (67), or so far unknown REE-dependent enzymes and regulator proteins. The latter explanation is of particular interest, as two recent studies provide strong evidence that specific importers that can transport Ln^3+^ into the cytoplasm of methylotrophic bacteria do exist (41, 68).

Altogether, the current study demonstrates that the utilization of REEs can influence important physiological traits of *P. putida*, which could be highly beneficial in competitive environmental niches such as the rhizosphere. The previously reported fertilizing effect of REEs on different food crops could hence be partially the result of increased competitiveness of plant growth promoting organisms such as *P. putida* during root colonization. This hypothesis is further supported by a recent study, which found that Pseudomonads predominantly thrive on root exudates *in vivo* and are hence enriched in the rhizosphere of *Arabidopsis thaliana* (10). As such, it will be interesting to see what future research will add to the currently emerging theme of REEs being an important micronutrient for methylotrophic and non-methylotrophic organisms.

---

**Table S1:**
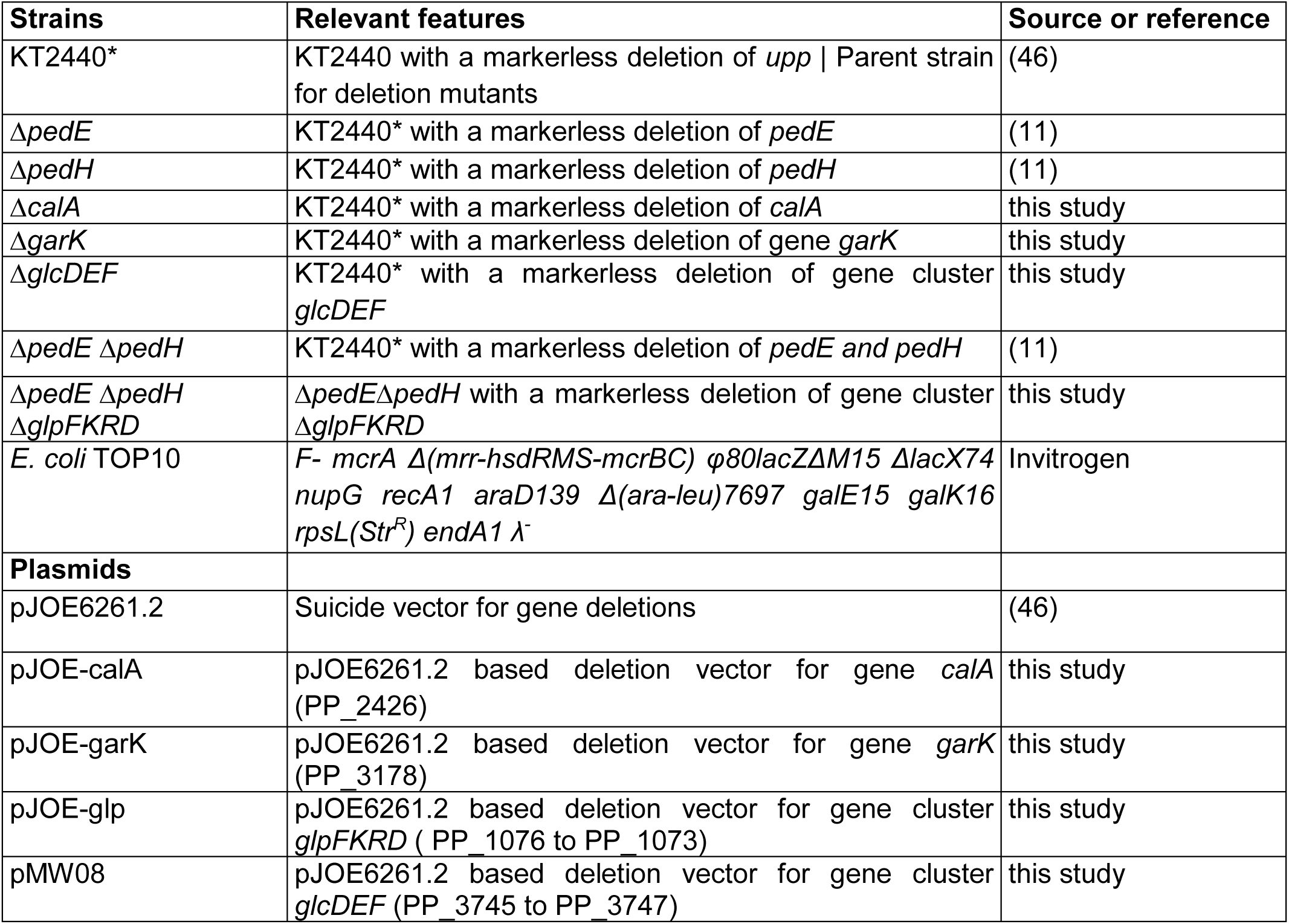
Strains and plasmids used in the study.

**Table S2:**
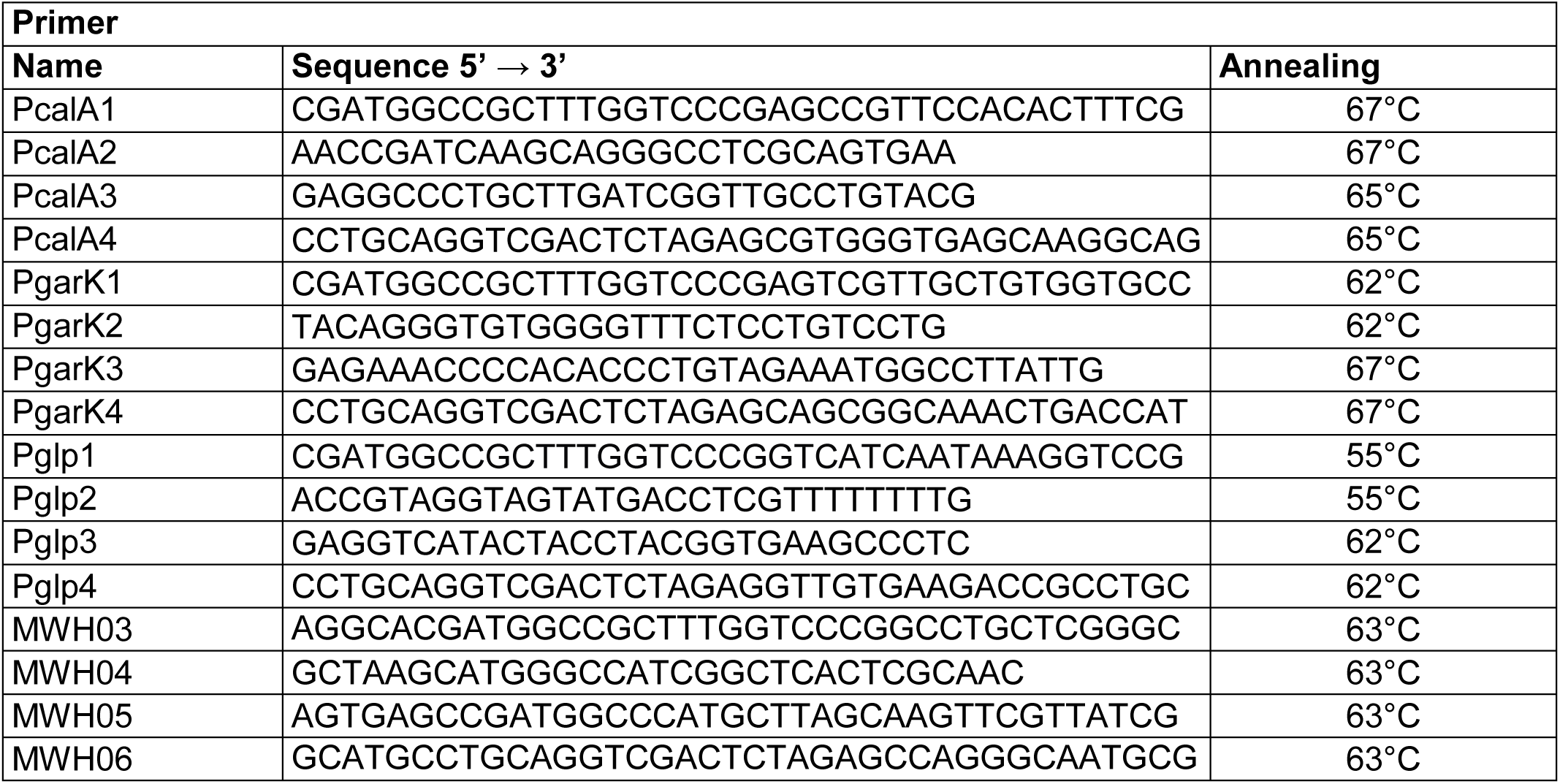
Primers used in the study.

**Table S3:** List of regulated proteins in presence of 10 μM La^3+^ compared to the absence of La^3+^ when grown with 2-phenylethanol as sole C-source.

| Locus Tag | Protein name | Fold change induction<br>(log<sub>2</sub>) | - log<sub>10</sub>(p-value) |
| --- | --- | --- | --- |
| PP_2679 | PedH | 3.41 | 2.48 |
| PP_3357 | Vdh | 2.65 | 2.92 |
| PP_5125 | MutM | 2.24 | 3.15 |
| PP_4905 | MotA | 1.84 | 2.01 |
| PP_0091 |  | 1.71 | 2.26 |
| PP_5157 |  | 1.59 | 2.20 |
| PP_0342 | WaaC | 1.43 | 2.46 |
| PP_3722 | Alr | 1.04 | 2.40 |
| PP_2674 | PedE | -1.49 | 5.90 |
| PP_2673 |  | -2.41 | 3.91 |

**Table S4:**
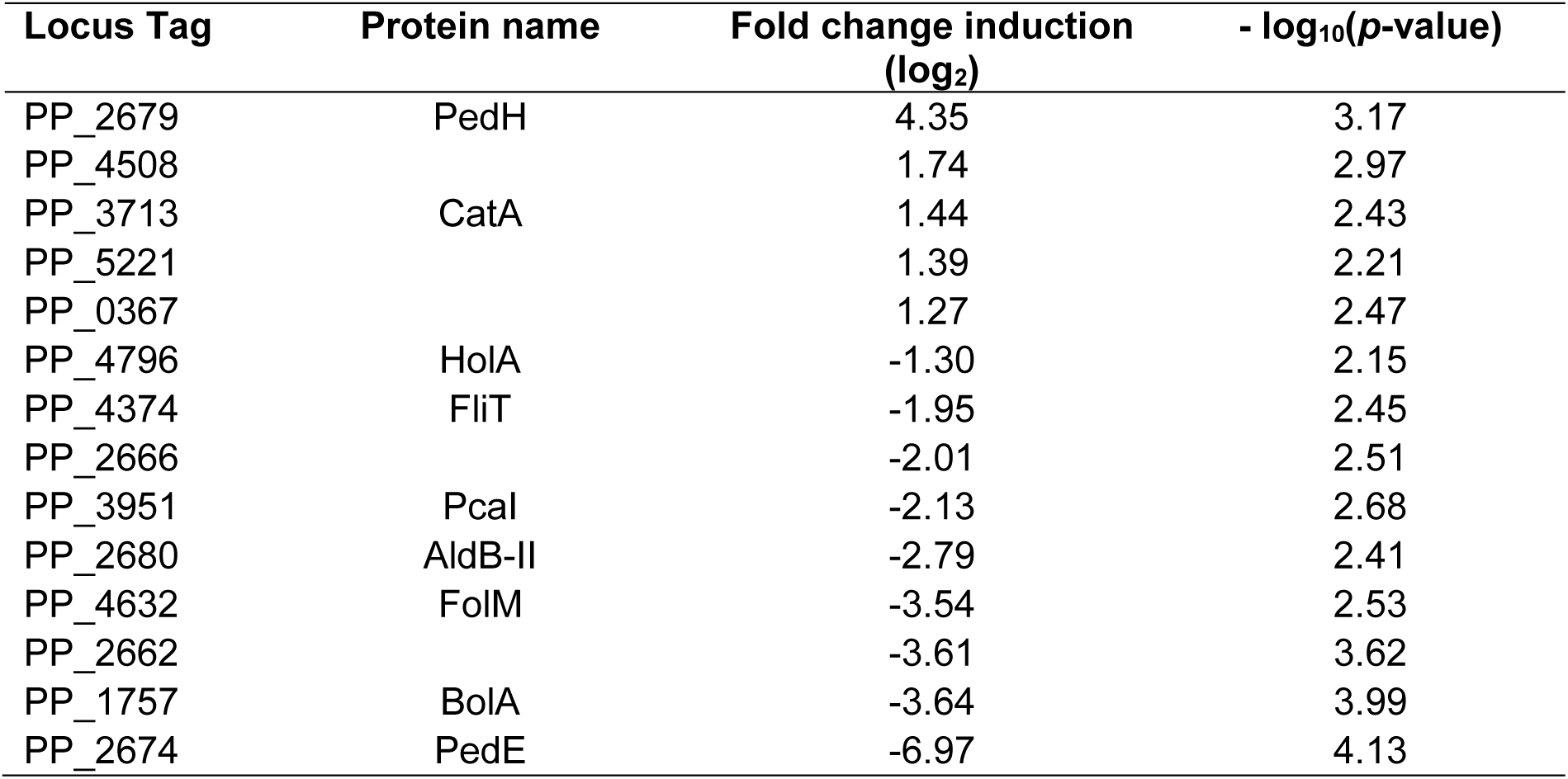
List of regulated proteins in presence of 10 μM La^3+^ compared to the absence of La^3+^ when grown with glucose as sole C-source.

**Table S5:** List of regulated proteins in presence of 10 μM La^3+^ compared to the absence of La^3+^ when grown with citrate as sole C-source.

| Locus Tag | Protein name | Fold change induction<br>(log<sub>2</sub>) | - log<sub>10</sub>(p-value) |
| --- | --- | --- | --- |
| PP_2491 |  | 2.27 | 4.32 |
| PP_0365 | BioC | 1.12 | 2.06 |
| PP_4157 | KdpE | -1.41 | 2.72 |
| PP_2674 | PedE | -2.56 | 2.80 |
| PP_0634 | PilA | -3.08 | 3.88 |

**Figure S1:**
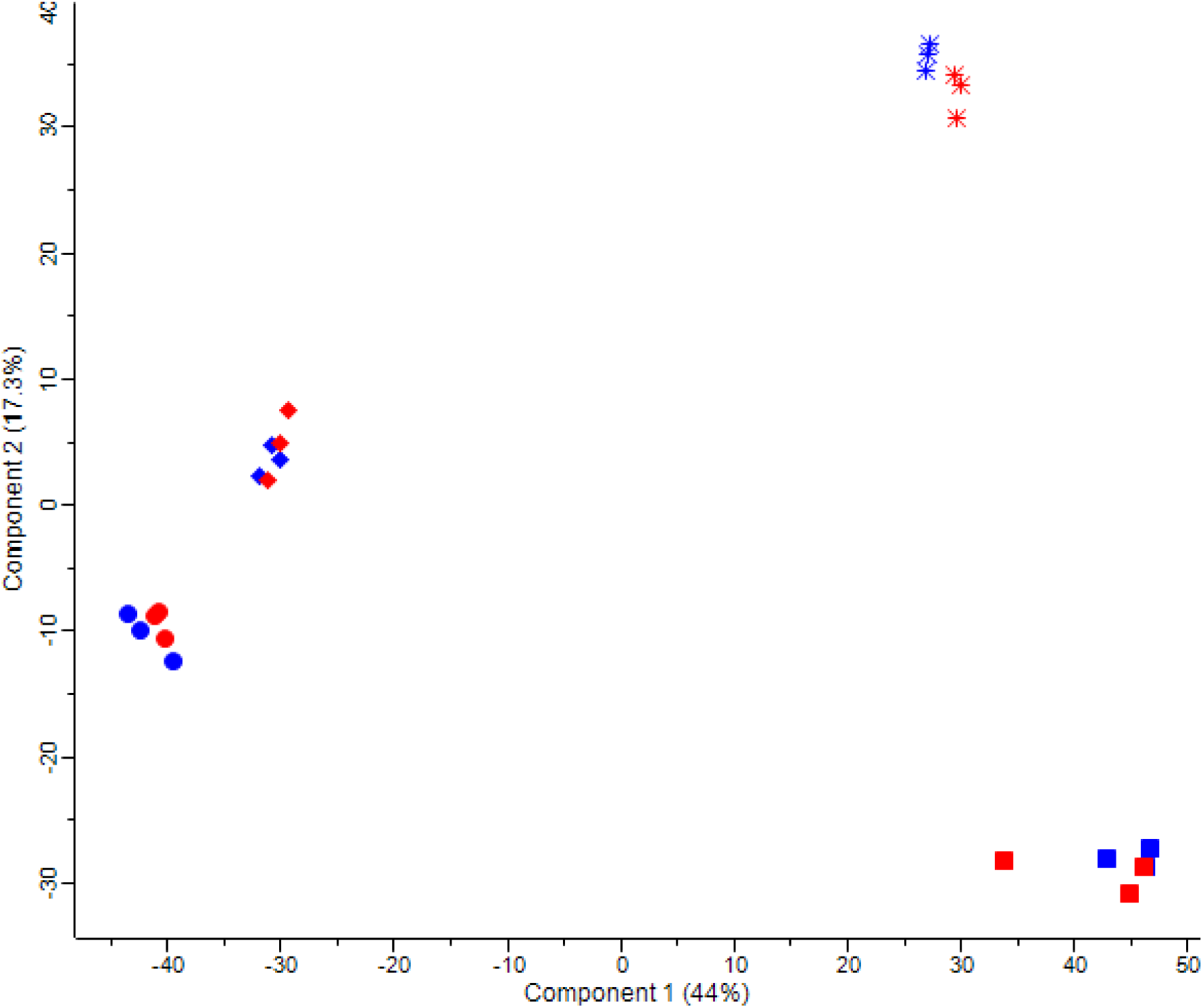
PCA comparing the four different carbon sources with and without La^3+^. Different carbon sources are indicated by squares (2-phenylethanol), circles (citrate), diamonds (glucose) and stars (glycerol). Samples with 10 μM La^3+^ or without La^3+^ in the medium are shown in blue and red, respectively. Biological replicates are indicated in the same colour. Samples can be separated according to different carbon sources while treatment with La^3+^ only showed minor effects.

